# Engineering Transcriptional Regulator Effector Specificity using Computational Design and In Vitro Rapid Prototyping: Developing a Vanillin Sensor

**DOI:** 10.1101/015438

**Authors:** Emmanuel L. C. de los Santos, Joseph T. Meyerowitz, Stephen L. Mayo, Richard M. Murray

## Abstract

The pursuit of circuits and metabolic pathways of increasing complexity and robustness in synthetic biology will require engineering new regulatory tools. Feedback control based on relevant molecules, including toxic intermediates and environmental signals, would enable genetic circuits to react appropriately to changing conditions. In this work, variants of qacR, a tetR family repressor, were generated by computational protein design and screened in a cell-free transcription-translation (TX-TL) system for responsiveness to a new targeted effector. The modified repressors target vanillin, a growth-inhibiting small molecule found in lignocellulosic hydrolysates and other industrial processes. Promising candidates from the *in vitro* screen were further characterized *in vitro* and *in vivo* in a gene circuit. The screen yielded two qacR mutants that respond to vanillin both *in vitro* and *in vivo*. We believe this process, a combination of the generation of variants coupled with *in vitro* screening, can serve as a framework for designing new sensors for other target compounds.

## Introduction

Engineering cells that contain circuits and novel metabolic pathways of increasing complexity and robustness in synthetic biology will require more sophisticated regulatory tools. The utility of a synthetic genetic circuits for real world applications is dependent on the ability to effectively trigger the circuit. While we can control the expression of target genes with transcriptional regulators, triggers for these transcriptional regulators are limited to a small number of molecules and other inputs (e.g. light) (*1*). As a consequence, most synthetic circuits right now are limited to proof-of-principle demonstrations without being extendable to real world applications. Feedback control based on relevant molecules, including toxic intermediates and environmental signals, would enable genetic circuits to react appropriately to changing conditions. This requires us to be able to transmit the levels of the relevant molecules to existing transcriptional control machinery. This work develops a framework to use a combination of sequence generation by computational protein design (CPD) and rapid prototyping using a cell-free transcription-translation (TX-TL) system to switch effector specificity of existing transcriptional regulators to respond to targeted small molecules of interest (Figure 1).

**Figure 1:**
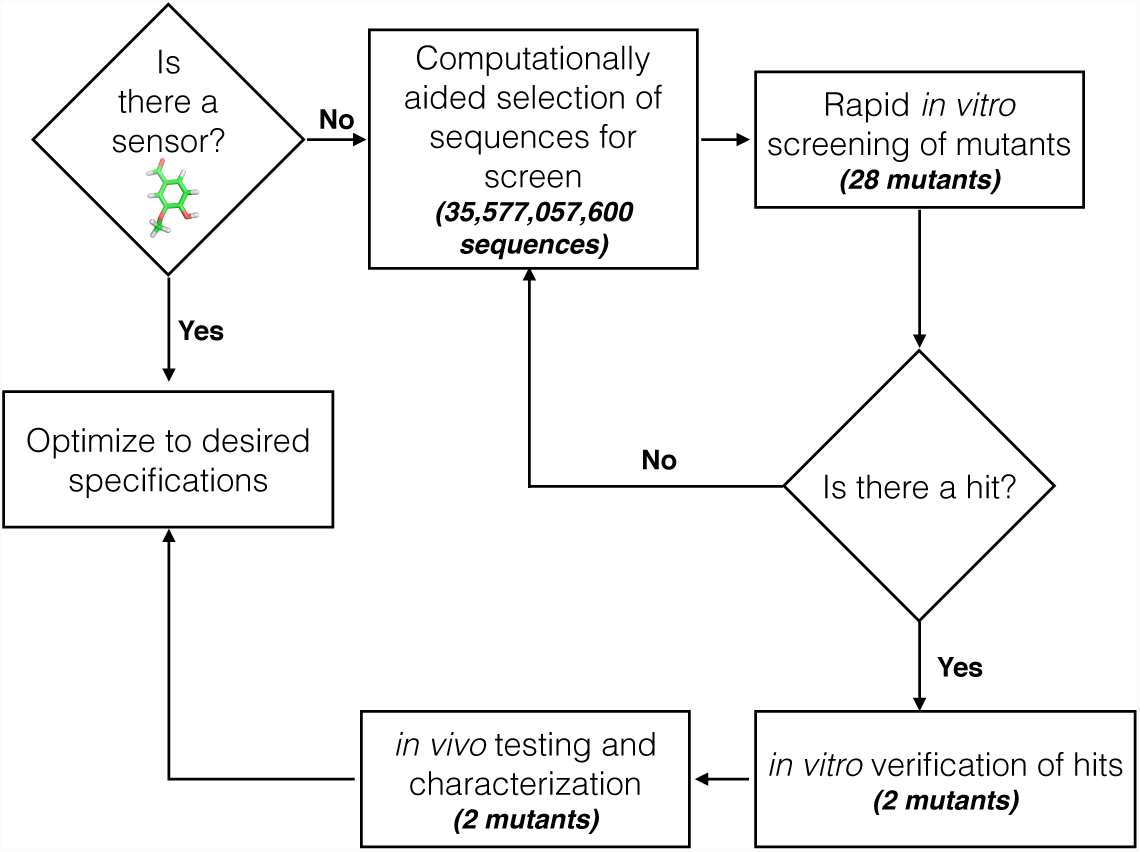
Workflow for generating novel sensors. The *in vitro* TX-TL platform allows for the rapid screening of sequences selected with the help of computational protein design. Hits from the *in vitro* screen are then verified by further *in vitro* testing. *In vivo* testing and characterization can then be performed to see if they meet the desired specifications. Further refinement of the hits through directed evolution or further computational design can be performed until specifications necessary are achieved. Numbers in parenthesis are the number of sequences considered by the computational algorithm, or the number of mutants assayed at the specified step for vanillin.

The tetR family is a large family of transcriptional regulators found in bacteria. They are named after the tetR repressor, which controls the expression of tetA, an efflux pump for tetracycline (*2*). They contain two domains, a helical-bundle ligand-binding domain and a helix-turn-helix DNA-binding domain. In the absence of their inducing molecule, tetR repressors bind to DNA, preventing the transcription of downstream genes. Inducer binding to the ligand-binding domain causes a conformational change in the DNA binding domain that causes dissociation from the DNA, allowing transcription of downstream genes. The tetR transcriptional regulation machinery has been used in the design of synthetic circuits, including the repressilator (*3*) and the toggle switch (*4*).

QacR is a tetR-family repressor found in *S. aureus* that controls the transcription of qacA, an efflux pump that confers resistance to a large number of quaternary anionic compounds. The protein has been studied because it is induced by a broad range of structurally dissimilar compounds (*5*). Structural examination of qacR in complex with different small molecules has shown that qacR has two different binding regions inside a large binding pocket. While qacR has multiple binding modes for various inducers, in all cases for which there are structures, binding of the inducer causes a tyrosine expulsion that moves one of the helices and alters the conformation of the DNA binding domain, rendering qacR unable to bind DNA (*6*–*8*). Crystal structures of inducer-bound forms of qacR and the qacR-DNA complex coupled with a definitive structural mechanism for qacR induction make it the ideal starting point for CPD of new transcriptional regulators. In this work, we describe our efforts to apply our framework to engineer qacR to sense vanillin, a phenolic growth inhibitor that is a byproduct of lignin degradation performed during the processing of biomass into intermediate feedstock in biofuel production (*9*).

## Results and Discussion

### Computationally Aided Selection of QacR Mutant Sequences

We created a computational model of vanillin to place into a crystal structure of QacR. Phoenix Match (*10*), a computational protein design algorithm was used to find potential vanillin binding sites close to the location of the tyrosine expulsion in the binding pocket of qacR (Figure 2A-B) while being in the proximity of amino acid positions that allowed for favorable pi-stacking and hydrogen bonding interactions. We used targeted ligand placement (*10*) to find potential binding positions for vanillin by defining an idealized binding site for the molecule. The algorithm yielded four potential binding positions for vanillin (Figure 2C). Computational protein sequence design was then used to select amino acid residues at positions around the potential vanillin binding sites. In order to minimize the possibility of steric clashes in the protein, we also performed calculations that considered both the DNA-bound state and the ligand-bound state using a multi-state design algorithm (*11*). Finally, we also ran calculations that included an energy bias to favor the wild-type residue. The lowest energy sequences from these four calculations (single-state biased, single-state non-biased, multi-state biased, and multi-state non-biased) were analyzed, and used as a guide to compile a set of ten mutants (Table S1) for *in vitro* testing. A more detailed description of the computational methods used can be found in the Materials and Methods section.

**Figure 2:**
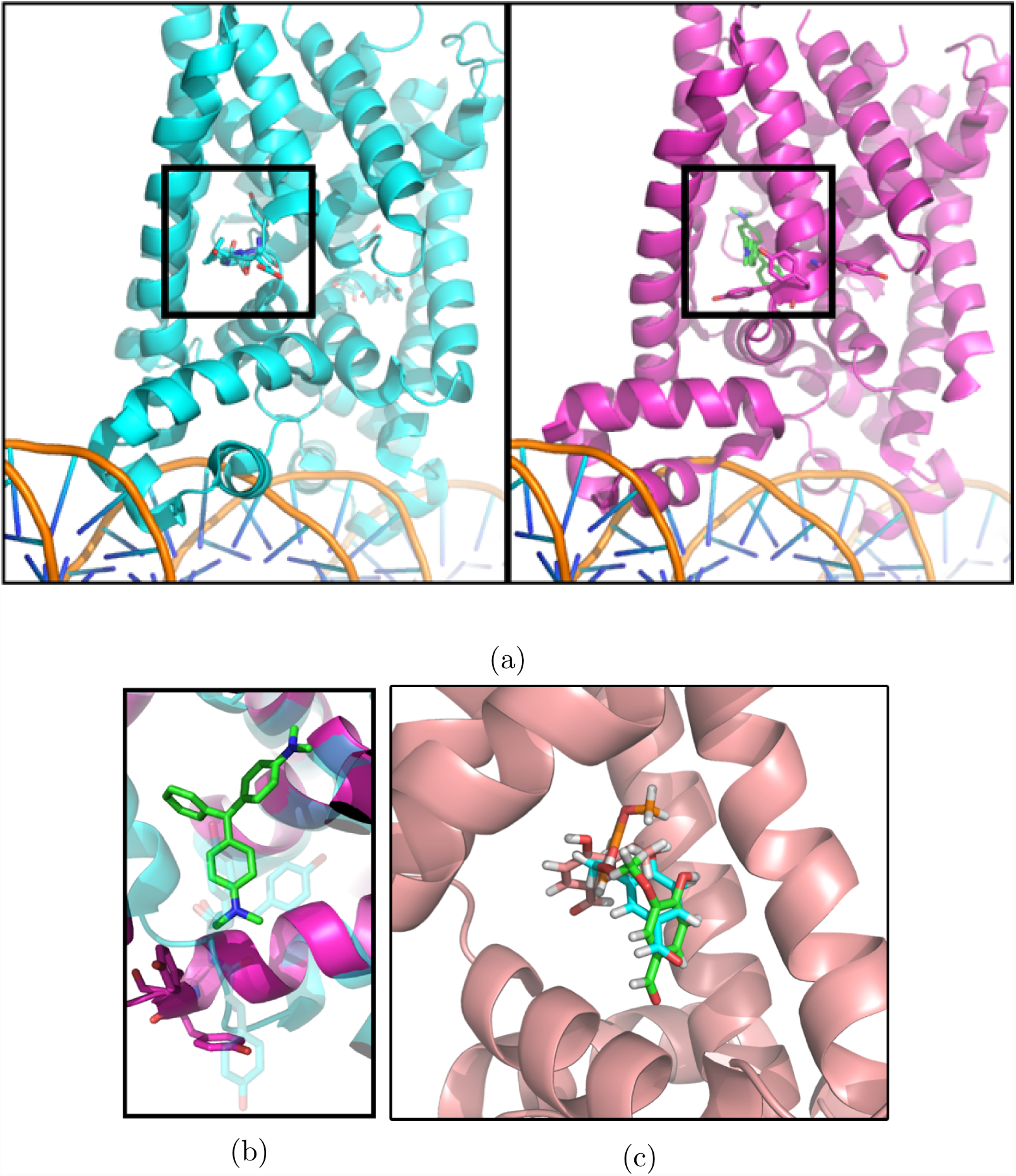
Computationally aided selection of qacR mutants. **(a)** PDB structures of the non-ligand bound (cyan, PDB ID: 1JT0) and ligand bound (magenta, PDB ID: 3BQZ) conformations of qacR. A conformational shift in the binding pocket occurs upon entry of the small molecule causing the protein to dissociate from DNA. **(b)** A closer look at the binding pocket of qacR, the binding of the ligand in green causes the displacement of three tyrosine shown as sticks in cyan and magenta. **(c)** Computational model for potential vanillin binding sites. Vanillin is shown as a different color in each of the four sites. A protein design algorithm was asked to suggest mutations for amino acids close to the potential binding sites to support the placement of vanillin in these sites.

### *In Vitro* Screening of Generated Sequences

We first decided to validate function of the wild-type protein. This was done by placing green fluorescent protein (GFP) downstream of the qacA promoter sequence (P_QacA_). While we observed a hundred-fold decrease in fluorescence in cells containing plasmids encoding the wild-type qacR gene in addition to P_QacA_–GFP, addition of berberine, a native qacR inducer, yielded no observable difference in fluorescence (Figure S1). We hypothesized that the inducer was not getting into the cells due to the differences in cell wall permeability between gram-positive and gram-negative bacteria. Because of this, we decided to use an *in vitro* transcription-translation (TX-TL) system to test the mutants (*12*).

The TX-TL system contains whole cell lysate from BL21 *E. coli* Rosetta 2, with no endogenous mRNA or DNA. A TX-TL reaction is typically done in a 10*μ*L reaction volume and contains the cell extract, an energy solution consisting of amino acids, nucleotides and 3-PGA, and DNA. It contains the transcriptional and translational machinery of *E. coli* allowing you to express proteins by adding plasmid DNA encoding genes you want expressed (Figure 3). Protein concentration can be controlled directly by varying the amount of DNA placed in the reaction. You can execute genetic networks in a TX-TL reaction by adding plasmids that contain proteins that interact. The TX-TL prototyping provides advantages over *in vivo* circuit testing as it allows us to control protein levels without worrying about promoter and ribosomal binding site strength. Cell wall permeability and protein toxicity are also not issues in the TX-TL system.

**Figure 3:**
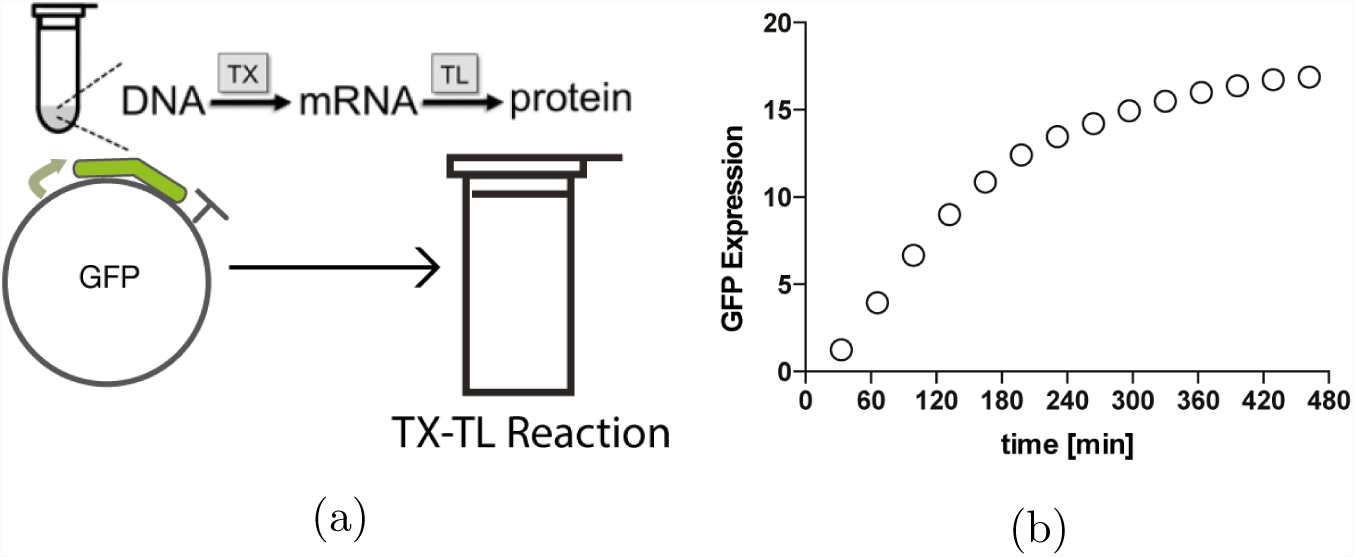
TX-TL allows us to prototype circuits *in vitro*. (a) TX-TL contains the transcriptional and translational machinery allowing you to express proteins in the reaction. Adding plasmid DNA encoding proteins allows for their expression and detection in TX-TL. deGFP expression from a TX-TL reaction with plasmid encoding GFP.

We first tested the wild-type protein in our TX-TL system. We observed an increase in GFP fluorescence as we increased the concentration of plasmid encoding P_QacA_–GFP from 2 nM to 8 nM (Figure 4A). The addition of plasmid encoding the qacR repressor to the system resulted in a decrease in fluorescence. Because of the high autofluorescence of berberine, we used dequalinium, a colorless native qacR inducer. The addition of dequalinium resulted in an increase in fluorescence until about 85% of the fluorescence when no DNA encoding repressor was present Figure 4B). These results demonstrated a functional wild-type qacR repressor in TX-TL. After validating the function of wild-type protein in TX-TL, we used the system to look at the functionality of the qacR mutants.

**Figure 4:**
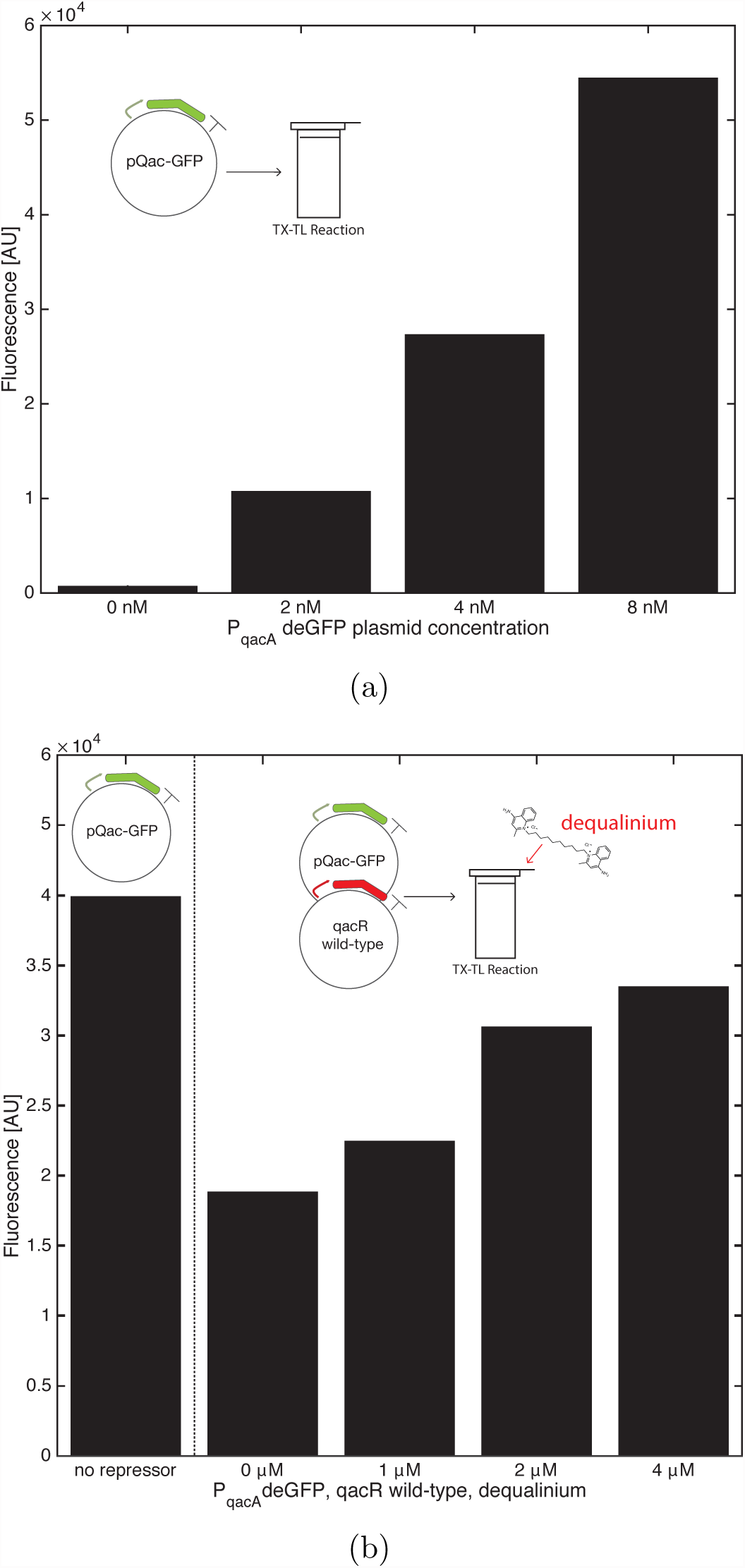
Validation of TX-TL screening. (a) GFP signal after three hours of a TX-TL reaction. Plasmid encoding GFP downstream of the native qac promoter was added to the TX-TL platform. Higher concentrations of plasmid yielded more GFP signal. (b) Response of wild-type qacR to dequalinium. DNA encoding GFP and wild-type qacR was added to the TX-TL system. Increasing fluorescent signal is observed with increasing concentrations of dequalinium. The highest fluorescent signal is observed when there is no repressor in the system, demonstrating the ability of TX-TL to test for qacR repression and de-repression.

None of the initial mutants showed any repression of GFP fluorescence. We analyzed the ligand bound and DNA-bound computational models of one of the qacR mutants that contained only three amino acid substitutions from a qacR mutant that was previously shown to be functional by Peters *et al.* (*8*). The computational model showed the potential for some mutations to cause steric clashes in the DNA bound state (Figure S2). We created a second library reverting either the 50th and 54th positions (A50F/W54L) or the 119th position (Y119L) to their wild-type identity (Table S2)

In order to determine if any of the mutants of our library warranted further characterization, we performed a rapid screen of 17 qacR mutants in TX-TL (Figure S3). Plasmids containing DNA that encoded each of the qacR variants or the wild-type qacR sequence were placed into a TX-TL reaction containing either water, dequalinium or vanillin. QacR activity was monitored by a plasmid encoding GFP downstream of P_QacA_. Two of the mutants, qacR2 and qacR5, displayed an increase in fluorescence in the presence of vanillin and dequalinium over water (Figure 5). We focused on these two mutants for further *in vitro* and *in vivo* characterization.

**Figure 5:**
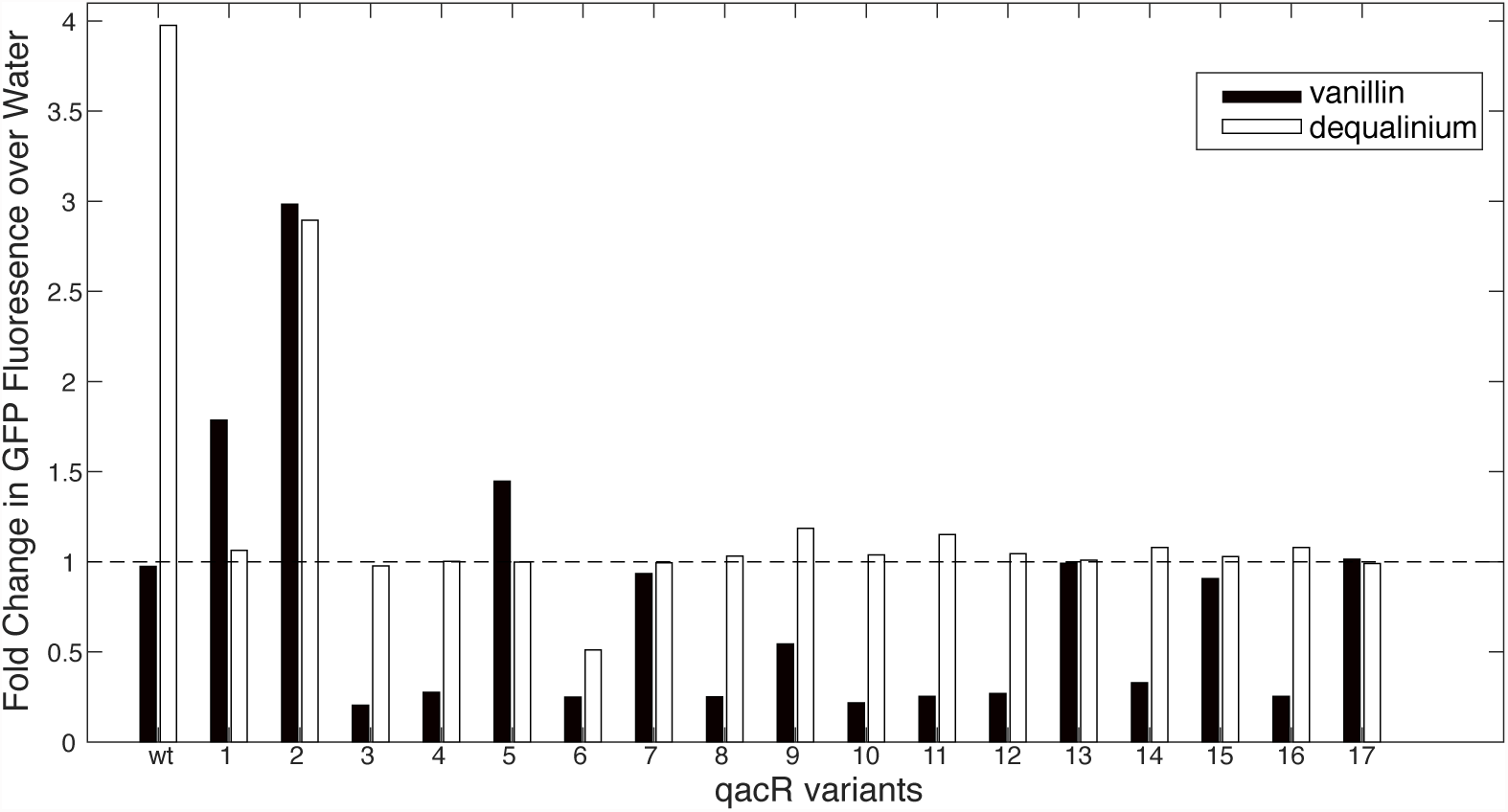
*In vitro* TX-TL screen of qacR mutants found potential candidates for further testing. Fold change in maximum fluorescence between water and inducer for qacR mutants. Seventeen qacR mutants were screened using TX-TL. Plasmids containing DNA encoding each of the qacR variants were placed into the system along with water, dequalinium (native qacR inducer) and vanillin. To monitor qacR response, a plasmid encoding GFP downstream of the native qacA promoter was also added to the system. qacR2 and qacR5 were selected for further characterization. qacR1 was not selected due to low signal (Figure S3)

### Further *In Vitro* Testing of QacR2 and QacR5

In order to verify the response of qacR2 and qacR5 to vanillin, we performed more extensive TX-TL tests on the mutants. TX-TL reactions were set up with a constant amount of reporter (P_QacA_–deGFP) plasmid and either no repressor (water), or plasmids encoding wild-type qacR, qacR2, or qacR5. Reactions were incubated for 85 minutes at 29 degrees Celsius to produce the repressor protein. This bulk reaction was then added to solution containing dequalinium, vanillin, or water. We monitored the rate of GFP production between the first and third hours of the reaction, where the rate of protein production appeared linear.

Figure 6 shows the ratio of GFP fluorescence between the case where there is no repressor, and each of the repressors tested with the different inducers. The wild-type qacR is able to inhibit the production of fluorescence to around 15% of its maximum value. The mutants are less efficient at repressing the production of GFP. Three times and four times more repressor DNA was added to the reactions of qacR2 and qacR5 respectively. In spite of the additional DNA, we do not observe the same level of repression that we see with the wild-type protein. Wild-type qacR is well induced by the native inducer, and we observed full derepression at the dequalinium concentration used. Induction of qacR2 and qacR5 with dequalinium is also observed, although to a lesser degree than the wild-type protein. QacR2 and qacR5 display a response to vanillin at the concentration we tested, while no response to vanillin was detected for the wild-type protein. The mutations introduced to the protein decrease the ability of the mutants to repress DNA. This could be due to protein instability, or due to a weaker protein-DNA interaction. However, these mutations also increase the sensitivity of the mutants to vanillin, allowing their response to be detectable in our *in vitro* platform.

**Figure 6:**
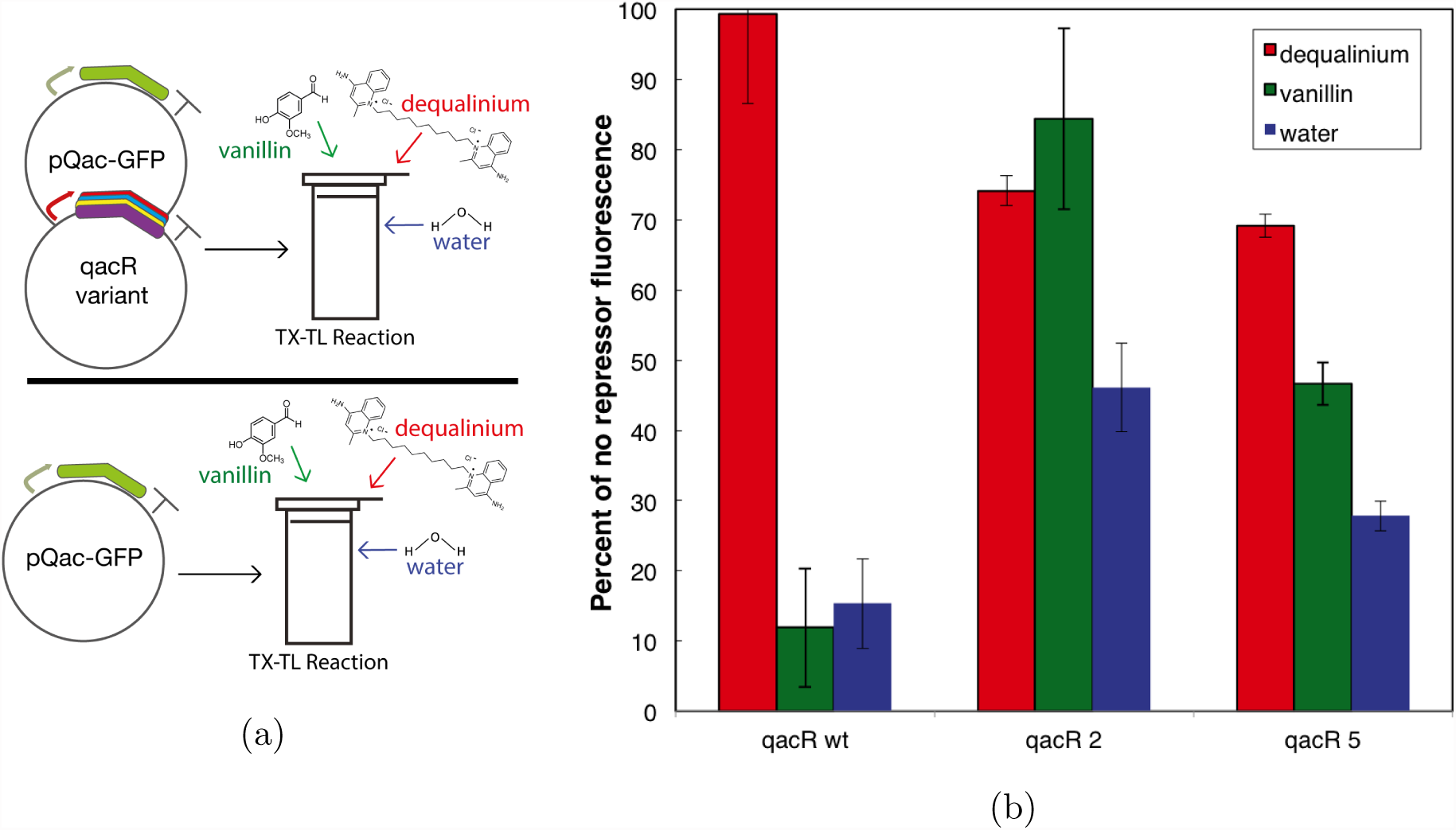
*In vitro* testing of qacR2 and qacR5. (a) TX-TL reactions were set up with plasmids containing GFP downstream of a qacR sensitive promoter in the presence and absence of plasmids containing the qacR variants under different inducer conditions. To account for inducer toxicity to the TX-TL reaction, and resource limitations from the production of the qacR repressor we normalized each condition to the reactions that only contained the GFP plasmid under different inducer conditions. (b) Ratio of the rate of GFP production between TX-TL reactions with and without repressor DNA. 10 *μ*M of dequalinium and 5 mM of vanillin was used to induce the production of GFP for each of the qacR variants tested.

We assumed that the maximum amount of GFP fluorescence that can be achieved for a specific inducer condition was when there is no repressor present. This takes into account potential toxicity of the inducer to the TX-TL reaction. The factors that can affect the ability of the particular repressor to reach this the no repressor case are resource limitations due to additional load from the production of the repressor DNA, and response of the repressor to the inducer in the reaction. We expect that resource limitations would have a negative effect on the ability of the repressor to reach the maximum fluorescence level. Conversely, response to repressor should have a positive effect in reaching the maximum fluorescence level.

### *In Vivo* Testing of QacR2 and QacR5

In order to further characterize the qacR mutants, and to see if we could detect vanillin in a more complex system, we decided to test the *in vivo* response of the qacR variants to vanillin. Plasmids containing genes that encode the wild-type qacR sequence, qacR2 or qacR5 downstream of P_Tet_ and GFP downstream of P_QacA_ were cloned into DH5*α*Z1 cells (Figure 7). For each of the qacR variants, we compared differences in fluorescence signal across increasing vanillin concentrations. We tested different repressor concentrations by varying the amount of anhydrous tetracycline (aTc) in the system. Similar to the *in vitro* experiments, and in order to get an idea for the maximum fluorescence the system could achieve, we grew cells that only contained GFP downstream of P_QacA_ without any repressor. Cells that were grown in higher aTc concentrations had a lower measured optical density (OD), indicating a slower doubling time. We hypothesize that this is due to the toxicity of the qacR repressor to the *E. coli* strain. Since qacR is not a native protein, it is possible that qacR is binding to locations in the *E. coli* genome. Interestingly, the differences in optical density measurements become less pronounced with increasing vanillin concentration, suggesting that vanillin may provide a mitigating effect to this toxicity. In order to account for differences in OD, fluorescence measurements were normalized to OD.

**Figure 7:**
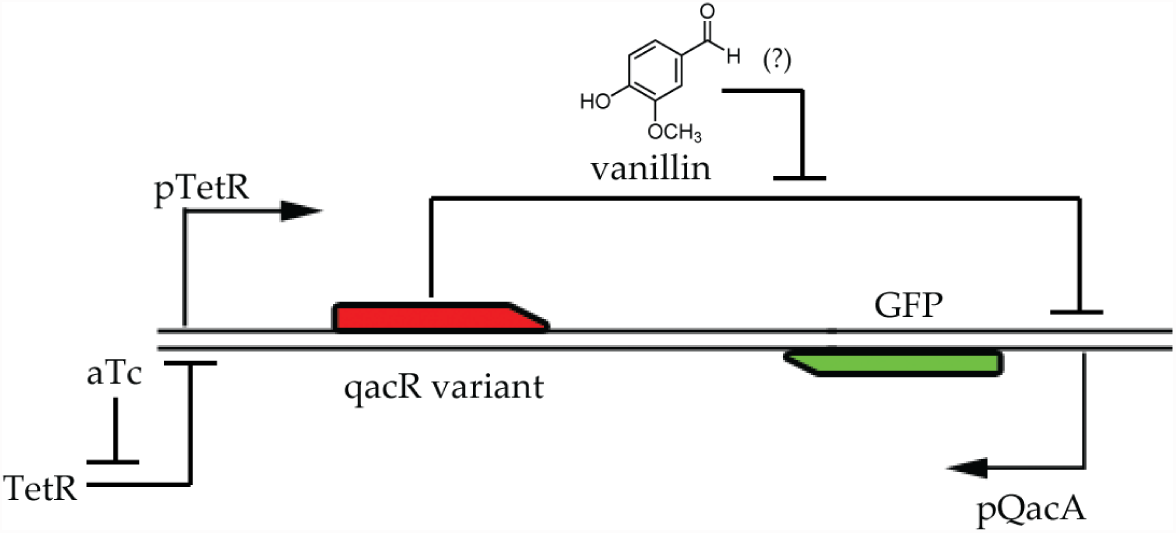
Circuit layout for *in vivo* tests. Genes encoding GFP under the control of the native qac promoter, and our QacR designs under the control of a tet-inducible promoter were placed in a single plasmid and transformed into DH5*α*Z1 cells. qacR levels were controlled using aTc for varying vanillin concentrations. Candidate designs that are responsive to vanillin should show an increase in fluorescence with increasing vanillin concentrations

The lowest OD measurements were observed for cells encoding the wild-type qacR at 12 ng/mL aTc where very little growth was observed for cells expressing the wild-type protein. At this aTc concentration, all of the cells expressing repressor exhibited lower optical densities when compared to cells that were only expressing fluorescent protein. The differences in optical density are less pronounced at lower aTc concentrations. When no aTc is present in the system, cells at the higher vanillin concentrations had lower ODs. At higher aTc concentrations, cells at higher vanillin concentrations had higher ODs. This implies that both the vanillin concentration and the expression of the repressor have an effect on cellular growth. The optical densities for the cells at different aTc and vanillin concentrations are shown in Tables S3-S6.

Figure 8A shows the effect of increasing the aTc concentration on the fluorescence of cells in the absence of vanillin. Similar to the *in vitro* tests, fluorescence was normalized to the no repressor case. Increasing the aTc concentration decreased the fluorescence of cells in the absence of vanillin, confirming that the qacR mutants are able to repress the expression of GFP at higher protein concentrations.

**Figure 8:**
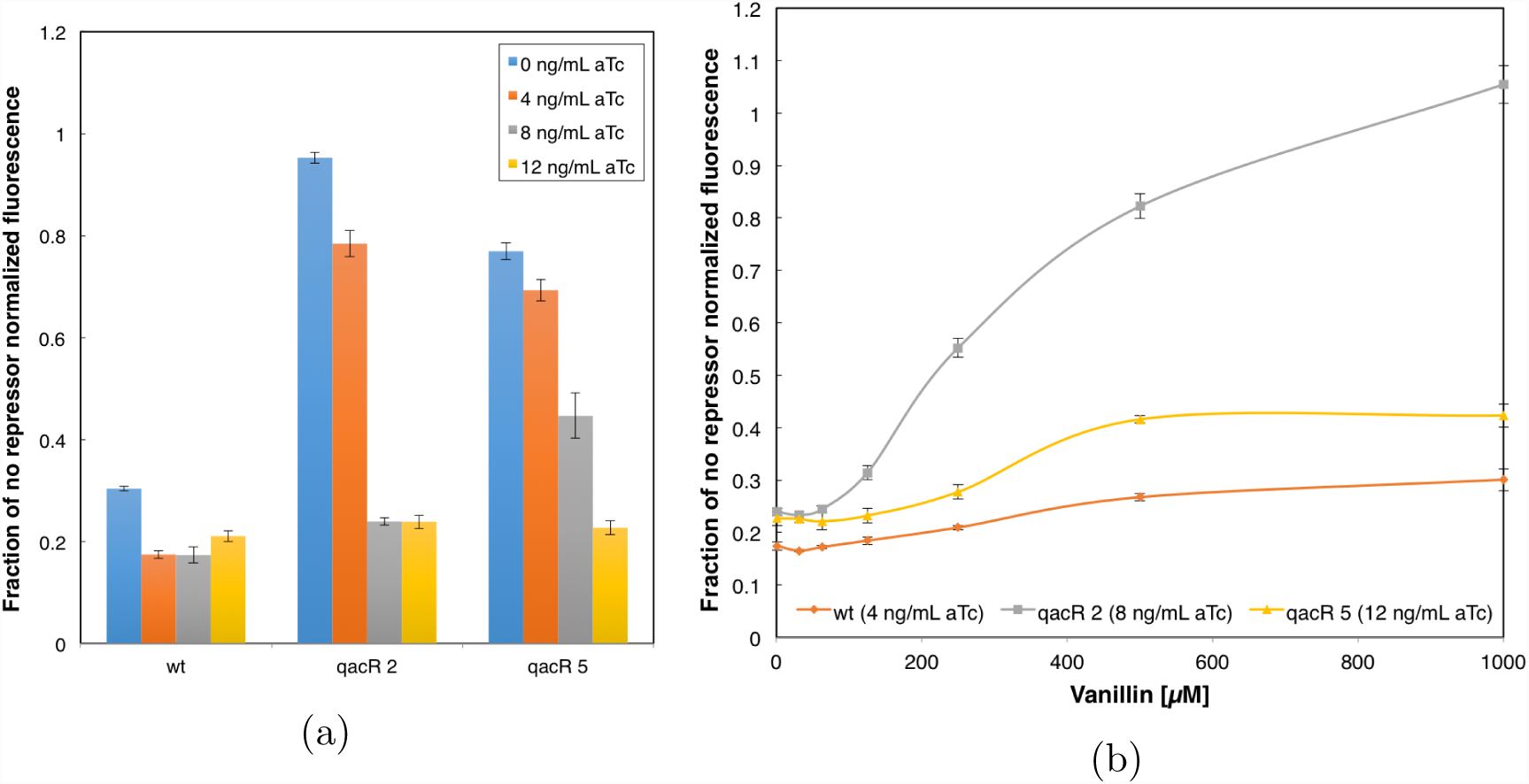
*In vivo* response of qacR to vanillin. Cells expressing GFP without any repressor were used as a control to normalize for differences in fluorescence due to aTc and vanillin levels. (a) All of the proteins are able to repress the expression of GFP. The wild-type protein is able to inhibit the expression of GFP at lower aTc concentrations, while higher aTc concentrations are necessary for the mutants to achieve a similar level of repression. (b) QacR mutants respond to vanillin in a concentration dependent manner.

The response of wild-type qacR, qacR2, and qacR5 to increasing vanillin concentrations is shown in Figure 8B. The response curves for each protein are plotted for the minimum aTc concentration such that maximum GFP repression is observed. This corresponds to aTc concentrations of 4, 8, and 12 ng/mL for wild-type qacR, qacR2, and qacR5 respectively. This is consistent with the *in vitro* data that more qacR2 and qacR5 DNA was required to repress the expression of GFP. Similar to the *in vitro* tests, we expect the ability of the cell to reach the maximum fluorescence level to be dependent on its response to inducer, and toxicity from vanillin and qacR. Indeed, cells expressing the qacR mutants exhibited an increase in fluorescence with increasing vanillin levels demonstrating that they are capable of sensing vanillin. While all three proteins appear to be sensitive to vanillin, the mutants exhibit a marked increase in sensitivity to vanillin. QacR2 displays a response that goes from approximately 20 percent of the fluorescence of the cells not expressing any repressor to matching the fluorescence of the non-repressed cells at 1 mM vanillin. QacR5 saturates at around 40 percent of the fluorescence of the non-repressed cells. This correlates with the *in vitro* data that show qacR2 achieving close to the non-repressed fluorescence, with qacR5 less sensitive to vanillin (Figure 6). Figure S4 shows the vanillin dosage response of qacR wild-type, qacR2, and qacR5 for different aTc concentrations tested.

### Framework Enables Engineering of Sensors through Rational Reduction of Design Space

The framework developed—a combination of sequence generation using computationally-aided design, preliminary screening with TX-TL, and *in vitro* and *in vivo* validation—can be used for other small molecule targets potentially facilitating the design of more sensors in synthetic circuits. While it is possible that the computational model of vanillin binding was inaccurate, the computational design provided value in drastically reducing the number of sequences to test into a figure that was experimentally tractable. Without the computational design to reduce the size of the design space, we would not have had effective starting points to attempt the engineering of a vanillin sensor.

The use of the *in vitro* cell-free system in a preliminary screen provides many advantages. It allows the screening of more mutants in a shorter amount of time. The simpler system also reduces the number of variables to consider. Complicating factors such as cell membrane permeability and cell growth do not need to be considered during this part of the screen. Repressors whose native inducers cannot enter the target organism can be used as starting points with the cell-free system. Finally, we can use this framework to target molecules that are known to be toxic to cells and measure engineering results in a cell-free context.

As a result of this process, we now have functional vanillin sensors that can be used in a feedback circuit that dynamically responds to vanillin. QacR2 and qacR5 can be used as a starting point for a synthetic circuit that responds to vanillin concentrations. While we only tested the protein in *E. coli*, recent work has developed a process that facilitates the transfer of prokaryotic transcription factors into eukaryotic cells, increasing the flexibility of the molecules for use in metabolic engineering (*14*). By linking a vanillin sensor to the expression of a gene that can mitigate the toxic effect of vanillin, such as an efflux pump or an enzyme which converts vanillin to a less toxic molecule, we can design a dynamic feedback circuit and potentially improve metabolic yield. It remains to be seen whether the sensors developed will have the required dynamic range or sensitivity for a functional feedback circuit; however, if a better sensor is needed these proteins can be used as a starting point for directed evolution in order to obtain a sensor with the desired properties.

## Materials and Experimental Methods

### Computationally Aided Selection of Mutant Sequences

An *in silico* model of vanillin was constructed using the Schrödinger software suite. Partial charges for vanillin were computed using Optimization in Jaguar version 7.6 (*15*) using HF/6-311G** as the basis set. Vanillin rotamers were chosen by looking at the ideal angles for the carbon hybrid orbitals. A model of an idealized vanillin binding pocket was designed by looking at the protein data bank for proteins that bound small molecules similar to vanillin, specifically PDBID 2VSU. Models of vanillin in the qacR binding pocket were generated using the Phoenix Match algorithm (*10*).

Vanillin was built off a native tyrosine residue (Y123), the primary interaction considered for the algorithm was a hydrogen bonding interaction between the hydroxyl group of the tyrosine with the methoxy and hydroxyl groups of the vanillin. We modified the energy function to include an energy bias for potential pi-stacking interactions between vanillin and tyrosine, phenylalanine, or tryptophan residues. We also included an energy bias hydrogen-bonding interactions with the methoxy, hydroxyl, and aldehyde groups of vanillin with serine, threonine, tyrosine, glutamine, or asparagine residues. The Phoenix Match algorithm was asked to return potential vanillin binding locations that contained interaction with the native tyrosine, at least one pi-stacking interaction, and at least two other hydrogen bonding interactions. Solutions from the algorithm were grouped together and resulted in four potential spots for vanillin. These locations were used as vanillin “rotamers” for computational protein design.

Monte Carlo with simulated annealing (*16*) and FASTER (*17*) were used to sample conformational space. A backbone independent conformer library with a 1.0 Å resolution was used for the designed residues (*10*). Designed residues were chosen by compiling a list of amino acid residues within 15 Å of vanillin. Table S7 shows the amino acid design positions, and the allowed amino acid residues for each position. Allowed amino residues for each site were selected by visually inspecting the qacR cyrstal structure with the potential vanillin binding locations. Rotamer optimization was allowed for other residues in the 15 Å shell in which mutations were not allowed. Computational models of qacR with vanillin present were scored using the PHOENIX forcefield with the inclusion of an additional geometry bias term that favored pi-stacking and hydrogen bonding interactions (*10*) that we used to find potential vanillin active sites. We considered solutions that both included and excluded *-*20 kcal/mol wild-type bias term in the energy function.

### Cell Free *In Vitro* Transcription-Translation System and Reactions

The transcription-translation reaction consists of crude cytoplasmic extract from BL21 Rosetta 2 *E. coli* (*12*). Preliminary tests were done with plasmids and inducers at the specified concentrations. For the initial screen, the qacR mutants were downstream of a T7 promoter. TX-TL reactions were run with 2 nM of the plasmid encoding the qacR variant, 0.1 nM plasmid encoding T7 RNA polymerase, and 8 nM plasmid encoding P_QacA_–deGFP. Vanillin was added at a concentration of 2.5 mM and dequalinium was added at 10 *μ*M.

For the *in vitro* tests to further characterize the hits, plasmids encoding qacR2 or qacR5 downstream of a tet-responsive promoter were used along with a plasmid encoding deGFP downstream of a qac-responsive promoter. Plasmids were prepared using the Macherey-Nagel NucleoBond Xtra Midi/Maxi Kit. Plasmid DNA was eluted in water and concentrated by vacufuge to the desired concentration. TX-TL reactions were set up as follows: 5 *μ*L of buffer, 2.5 *μ*L of cell extract and 1.5 *μ*L repressor DNA at a specific concentration was mixed and incubated at 29 C for 75 minutes to facilitate the production of repressor DNA. This mix was then added to a mixture of 1 *μ*L deGFP plasmid and 1 *μ*L of an inducer stock. Measurements were made in a Biotek plate reader at 3 minute intervals using excitation/emission wavelengths set at 485*/*525 nm. Stock repressor plasmid concentrations were 243 nM, 729 nM, and 972 nM for qacR wild-type, qacR2, and qacR5, respectively. The deGFP plasmid concentration was approximately 397 nM. Inducer concentrations were 5 mM for vanillin, and 10 *μ*M for dequalinium.

Experimental conditions were done in triplicate and the error bars are the error propagated from the standard deviation of the means.

### Cell Strain and Media

The circuit was implemented in the *E.coli* cell strain DH5*α*Z1, a variant of DH5*α* that contains a chromosomal integration of the Z1 cassette (*18*). The Z1 cassette constitutively expresses the TetR and LacI proteins. All cell culture was done in optically clear M9ca minimal media (Teknova M8010).

### Genes and Plasmids

DNA encoding the qacR genes was constructed using overlap extension PCR. Plasmids used contained chloramphenicol resistance with a p15a origin of replication.

### *In Vivo* Experiments

Cells were grown in at least two consecutive overnight cultures in M9ca minimal media. On the day of the experiment, overnight cultures were diluted 1:100 and grown for 5 hours to ensure that the cells were in log phase. Cells were then diluted 1:100 into fresh media at the specified experimental condition. Cells were grown in these conditions at 37C for 12–15 hours in Axygen 96 well plates while shaking at 1100 rpm. Endpoint fluorescence was measured by transferring the cells to clear bottomed 96-well microplates (PerkinElmer, ViewPlate, 6005182). GFP was read at 488/525 with gain 100.

Analysis of the data was done by taking fluorescence readings for each independent well. Experimental conditions of the qacR proteins were done in triplicate and repeats were averaged. Error bars shown are the error propagated originating from the standard deviation of the mean.

## Acknowledgement

The authors thank Jongmin Kim and Jackson Cahn for reading the manuscript. This research was conducted with supprot from the Institute for Collaborative Biotechnologies through grand W911NF-09-0001 from the U.S. Army Research Office. Additional support was granted in part by the Benjamin M. Rosen Bioengineering Center, the Gordon and Betty Moore Foundation through Grant GBMF2809 to the Caltech Programmable Molecular Technology Initiative, and DARPA through the Living Foundries Program.

